# Evaluation of near-infrared light therapy for the treatment of neurodegenerative diseases: Limited penetration depth into the brain likely hinders efficacy

**DOI:** 10.1101/2024.11.18.624091

**Authors:** Jessica Tittelmeier, Leon Kaub, Stefan Milz, Daniela Kugelmann, Patrick R. Hof, Christoph Schmitz, Carmen Nussbaum-Krammer

**Affiliations:** Department of Anatomy II, Faculty of Medicine, Ludwig-Maximilians-University (LMU) Munich, Pettenkoferstr. 11, 80336 Munich, Germany; Department of Anatomy I, Faculty of Medicine, Ludwig-Maximilians-University (LMU) Munich, Pettenkoferstr. 11, 80336 Munich, Germany; Nash Family Department of Neuroscience and Friedman Brain Institute, Icahn School of Medicine at Mount Sinai, 787 11th Avenue, New York, NY 10019, USA

**Keywords:** Neurodegenerative diseases, Alzheimer’s disease, Near-infrared light therapy, Transcranial photobiomodulation

## Abstract

**Background:** Near-infrared (NIR) light therapy is used to treat various musculoskeletal disorders. It has been proposed that transcranial NIR light treatment may also be beneficial for Alzheimer’s disease (AD). However, the ability of NIR light to penetrate the scalp and skull efficiently and induce cytoprotective responses in the brain parenchyma has not been sufficiently examined so far. This study aimed to evaluate whether the amount of NIR light that can penetrate through the human skull can cause a biological effect.

**Methods:** Three commercially available devices (a medical laser emitting light at a wavelength of 905 nm and two LED helmets operating at wavelengths of 810 nm and 1070 nm, respectively) were used to measure the NIR light transmittance through human post-mortem skulls with a thermal power sensor. Furthermore, the biological effects of the fraction of light power that passed through the skull were investigated in a human neuronal cell line and in *C. elegans*.

**Results:** The 905 nm laser achieved transmittances of up to 0.31% (173 µW/cm^2^) of its input power, and the LED helmets 0.71% (41 µW/cm^2^; 810 nm) and 0.45% (19 µW/cm^2^; 1070 nm) of their respective input powers. NIR light exposure at a power density of 134 mW/cm^2^ was sufficient to activate mitochondrial metabolism in cultured human neurons and *C. elegans*, as demonstrated by increased cytochrome c oxidase activity and induction of mitochondrial chaperones. However, this stimulatory effect was no longer observed when the applied power density was reduced to 2.5 mW/cm^2^.

**Conclusions:** More than 99% of the NIR light emitted by the investigated devices was either absorbed or scattered by the human skull. The residual NIR light that would reach underlying brain structures was too weak to elicit biological effects. In conclusion, NIR light treatment is unlikely to be effective to treat brain diseases such as AD due to the low penetrability of the skull.

## INTRODUCTION

Near-infrared (NIR) light has emerged as a popular method for the treatment of various conditions, particularly musculoskeletal disorders. It has been successfully used in physiotherapy and sports medicine to relieve pain and accelerate tissue regeneration.^1,2^ Additionally, NIR light therapy has shown promise in improving recovery from peripheral nerve injuries in animal studies, demonstrating that NIR light can decrease neurodegeneration and increase axonal growth and myelinization.^3^

While the underlying molecular and cellular mechanisms of NIR light therapy remain poorly understood, its therapeutic effects appear to be mediated primarily not by temperature but by photobiomodulation (PBM), in which the light energy is hypothesized to be absorbed by cellular components containing chromophores, leading to a range of beneficial effects.^4,5^ Absorbed photons have been shown to stimulate the enzyme cytochrome c oxidase (COX), the complex IV of the mitochondrial electron transport chain, upregulating mitochondrial respiration, resulting in increased production of reactive oxygen species (ROS) and adenosine triphosphate (ATP), which is essential for cellular energy supply.^4^ ROS trigger the redox-sensitive NFkB signaling pathway that regulates the expression of anti-apoptotic and pro-survival genes.^6^ The ability of NIR light to promote the phosphorylation of endothelial nitric oxide synthase (eNOS) leads to enhanced nitric oxide (NO) bioavailability in endothelial cells, modulating vasodilation.^7,8^ The resulting enhanced tissue perfusion improves the supply of oxygen and nutrients to the tissues and supports healing processes. NIR light also modulates the release of cytokines and other inflammatory mediators, reducing inflammation and pain and facilitating cellular repair.^5^

Growing evidence suggests that inflammatory processes and altered mitochondrial bioenergetics play crucial roles in the development and progression of Alzheimer’s disease (AD) and other neurodegenerative diseases,^9-11^ positioning NIR light as a potentially effective therapeutic strategy for these conditions. The non-invasive transcranial application of NIR light could also overcome the challenge posed by the blood-brain barrier, which limits the delivery of drugs into the brain. However, due to the limited number of treated patients and the absence of appropriate controls in corresponding clinical studies, definitive conclusions cannot yet be drawn regarding whether this treatment offers a genuine improvement over placebo.^12-14^

NIR light with wavelengths ranging from approximately 750 to 1100 nm can penetrate deeper into tissues compared to visible or ultraviolet light, reaching underlying structures without causing significant damage to the skin surface.^15^ However, we recently showed that approximately 90% of NIR light energy is absorbed within a depth of 5 to 10 mm in different animal tissue types, including skin, fat, muscle, tendon and bone tissue.^16,17^ This raises concerns about the efficiency of NIR light in passing through the different layers of the human head. Despite the broad applicability of NIR light therapy, its efficacy to penetrate the human skull and effectively stimulate brain cells has remained largely unexplored.

This study tested whether the amount of NIR light that can penetrate through the human skull is able to elicit a biological effect. Specifically, the extent to which NIR light emitted from three commercially available devices (a medical laser and two LED helmets) can penetrate the outer layers of the human head (skin, cranial bone, and dura mater) was measured using a thermal power sensor. In addition, the impact of the fraction of NIR light that can pass through the skull on mitochondrial activity was examined *in vitro* using human neurons and *in vivo* using *C. elegans* to evaluate the potential efficacy of NIR light therapy for neurodegenerative conditions.

## METHODS

### Transcranial NIR light transmittance measurements

Three human skulls (including shaved skin, subcutaneous tissue, cranial bone, and dura mater; see Supplementary Figure 1) were examined to account for the variability within the specimens. All skulls were formalin fixed and derived from bodies that had been donated to the Department of Anatomy I at LMU Munich. Experiments with cadaver skulls were performed in context of the body donor program of the Department of Anatomy I at LMU Munich. Body donors gave their written and informed consent for research purposes. The local ethical committee of LMU Munich gave their confirmation that no further approval for the experimental protocol is required. The skulls were stored at 4 °C and warmed up to room temperature before the measurements. Prior to measurements, the skulls were submerged in water for 24 hours to remove residual formalin.

To assess the extent to which NIR laser light can penetrate the human skull, three commercially available devices were used: a medical laser device emitting light with a wavelength of 905 nm (hereafter referred to as 905 nm laser; Dolorclast High Power Laser; Electro Medical Systems, Nyon, Switzerland) and two LED helmets with different wavelengths (810 nm and 1070 nm, hereafter referred to as 810/1070 nm LED helmets; Suyzeko Photobiomodulation Helmet, model GY-PDT1, Shenzhen Guangyang Zhongkang, Shenzhen, China).^18,19^

The 905 nm laser emitted laser pulses with a peak power of 300 W and a pulse length of 100 ns, resulting in an average power of 1.2 W at the maximum repetition rate of 40 kHz.^16,20^ For the measurements conducted with the 905 nm laser, the skulls were placed on a metal frame (Supplementary Figure 1). The focused nature of light emitted by the 905 nm laser allowed for the measurement of transmittance through the skulls at three distinct positions for frontal, parietal, and occipital sites (indicated in Supplementary Figure 1).

Each LED helmet was equipped with 256 NIR and four red light LEDs mounted on the inner side of the helmets and covered with acrylic glass. As stated by the manufacturer, both LED helmets were designed to emit pulsed NIR light. A temporal characterization of the light pulses was conducted with a fast photodiode sensor (FPD-VIS300; Ophir Spiricon Europe, Darmstadt, Germany) that was connected to an oscilloscope (MSO7024; Rigol Technologies, Suzhou, China). Temporal profiles were recorded at various repetition rates (1, 10 and 100 Hz) and maximum intensity. At higher repetition rates, interference signals from the multiple LEDs operated simultaneously precluded further recordings.

Preliminary measurements revealed significant scattering of light emitted by the LED helmets. When more than 75 LEDs were measured, a fraction of the IR light was observed to bypass the skulls, reach the sensor, and bias the recordings (data not shown). As a result, all areas of the LED helmets that did not directly illuminate the skull had to be shielded with aluminium foil to prevent the light from traveling past the outside of the skull. To evaluate the effect of individual LEDs, transmittance was measured with an increasing number of LEDs, up to the maximum of 75 LEDs (Supplementary Figure 1). To measure the light transmission, the skulls were placed inside the upside-down helmets (Supplementary Figure 1). The LED helmets were operated at their maximum repetition rate of 20 kHz.

Light power was measured with a thermal power sensor specifically designed for divergent incoming light (Model 3A-P-FS-12; Ophir Spiricon Europe) (Supplementary Figure 1). The sensor has a 12-mm diameter aperture, an absorber surface area of 9.6 × 9.6 mm, and a 6-µW power noise level. The sensor was equipped with a window blocking thermal radiation with a wavelength greater than 2100 nm. To measure light transmission, the NIR light sources were positioned on the outside of the skull and the power sensor was placed inside. The distance between light emitter and sensor was 15.7 cm for the 905 nm laser. For the LED helmets, the sensor was placed directly above the edge of the helmets. To prevent interference from external light, an aluminium shield was placed around the sensor aperture (Supplementary Figure 1). The measurements were conducted in a thermally stabilized, dark laboratory setting.

Recordings of the transmitted light were seven minutes in duration, with the light emitters being switched on and off in one-minute intervals (Supplementary Figure 2). This resulted in three intervals in which the devices were activated and four baseline measurements between each interval. Mean values were calculated from 40 seconds of data collected during each on- and off-time (Supplementary Figure 2). At each interval, a measurement was computed as the mean on-value minus the mean baseline, resulting in a total of five measurements for each recording. The initial baseline measurement was excluded due to thermal instabilities and sensor drift.

Light intensity was measured before and after passing through the skulls, allowing for the determination of both the absolute light power emitted from the devices and transmitted through the skull (in µW/cm^2^), as well as the calculation of the percentage of NIR light that penetrated the skull.

### Cell culture

SH-SY5Y human neuroblastoma cells (ATCC, Manassas, VA, USA) were cultured in DMEM containing high glucose, GlutaMAX Supplement, and pyruvate (Thermo Fisher Scientific), supplemented with 10% FBS (Thermo Fisher Scientific) at 37 °C and 5% CO2. Regular mycoplasma tests were performed (Eurofins Genomics, Ebersberg, Germany).

### COX activity assay in cells

COX activity in cells was assessed according to Zhang et al.^21^ Cells were seeded on Poly-l-lysine-coated coverslips. On the next day, the cells were exposed to the 905 nm laser at respectively 134 mW/cm^2^ or 2.5 mW/cm^2^ for 5 min and recovered for 3 hours. For cells treated with sodium azide (NaN3), 100 µM NaN3 (Sigma-Aldrich) were added to the cell culture media 24 hours before laser exposure to inhibit COX activity. Cells were then fixed in 4% paraformaldehyde in PBS for 10 min. After being washed three times with 10X TBS, COX reaction media consisting of 1% COX (Sigma-Aldrich), 1% 3,3′-diaminobenzidine (DAB) (Sigma-Aldrich) in 10X TBS was added to cells for 4 hours. COX activity was assessed by using DAB as electron donor (the enzyme oxidases DAB, which generates a brown, insoluble indamine polymer that is deposited at the reaction site). After being washed three times with 10X TBS, cells were mounted and imaged using an Axio Imager M2 (Zeiss, Jena, Germany) equipped with a 63× oil immersion objective and a Zeiss AxioCam MRc camera.

### Maintenance of *C. elegans* and age synchronization

Nematodes were cultured using standard methods.^22^ Animals were grown on nematode growth medium (NGM) plates seeded with *E. coli* strain OP50 at 20 °C. Age-synchronization of a population was achieved by bleaching. Briefly, gravid adults were dissolved in 20% sodium hypochlorite solution. The surviving *C. elegans* embryos were hatched overnight in M9 buffer with gentle rocking at 20 °C. The next day, L1 larvae were distributed onto plates and grown until they reached adulthood. The following strains were purchased from the Caenorhabditis Genetics Center (CGC, University of Minnesota, USA): Bristol N2 wild-type, SJ4100 (hsp-6p::GFP), SJ4058 (hsp-60p::GFP), and CF1553 (sod-3p::GFP). The strain CNK199 (HSP-16.2::mCherry) expressing endogenously tagged HSP-16.2 has been generated for and described in a previous study.^23^

### Laser exposure and quantification of fluorescence intensity

Adult day 1 worms were transferred to empty NGM plates and exposed to the 905 nm laser for 5 or 30 minutes. The distance between the laser handpiece and the NGM plates was set at 7.8 cm to ensure full illumination of the entire 5.5-cm plate. At this distance, the incoming power density of the 905 nm laser was 134 mW/cm^2^. Due to the divergence of the light beam and a larger distance between the laser handpiece and the sensor for the transmittance measurements on the human skulls, the power density of the 905 nm laser used here was larger than in the skull transmittance experiments. To reduce light power, a thin, white-colored polystyrene disc was placed under the laser, acting as a reflective neutral density filter. The polystyrene disc blocked the power density to 2.5 mW/cm^2^, equating to 1.8% of the original laser power density. After exposure, worms were recovered on seeded NGM plates and imaged 24 h post-exposure. Control worms were also placed on empty NGM plates for the equivalent durations in the dark and recovered on seeded NGM plates. For imaging, worms were mounted on agarose pads (8-10% agarose in M9 buffer) with a drop of mounting mix (2% w/v levamisole and 50% v/v nanosphere size standards; (Thermo Fisher Scientific, Waltham, MA, USA) and covered with a coverslip. Images were captured using a Leica M205 FA fluorescence stereomicroscope (Leica, Germany). Fluorescence intensity was quantified in FIJI (ImageJ, https://imagej.net/) by drawing a region of interest around the brightfield image of an animal and measuring the corresponding intensity in the fluorescence channel. Intensity was normalized to the control for each experiment.

### COX activity assay in *C. elegans*

Both, worms exposed to 905 nm NIR light and control worms were washed off NGM plates and transferred to 15-ml Falcon tubes containing 1X OP50 bacterial suspension. Tubes were placed on a rocker for a 3-hour recovery period. Following recovery, worms were pelleted by centrifugation at 300 x g for 1 minute and subsequently washed once with M9 buffer to remove OP50. Worms were then embedded for cryosectioning. The COX activity assay was performed according to Hench et al.^24^ Briefly, the worms were transferred into 1 ml of TissueTek^®^ carbowax tissue embedding medium (Sakura Fintek, Netherlands) in a 5-ml tube, then centrifuged at 1300 x g for 5 minutes, yielding a worm pellet, which was transferred into a Cryomold® (Sakura Fintek, Netherlands). To reduce stress-induced curling and promote sedimentation in longitudinal orientation, the cryo-molds were incubated at approximately 8 °C for 10 minutes. The blocks were then frozen in a glass dish filled with isopentane that had been pre-cooled in liquid nitrogen, thereby avoiding direct contact between liquid nitrogen and the sample. Frozen blocks were stored at −20 °C and sectioned on a CM1950 cryo-microtome (Leica, Germany) into 7-µm thick sections, which were mounted on SuperFrost® plus microscopy glass slides (Thermo Fisher Scientific). The sections were airdried for 2 hours and then washed twice in PBS (pH 7.4) for 10 minutes, followed by incubation at room temperature with a reaction solution containing 0.05% DAB, 0.1% cytochrome c (Sigma-Aldrich, St. Louis, MO, USA) and 83 mg/ml sucrose in PBS. The reactions were terminated after 90 minutes by washing the sections three times with PBS for 5 minutes each. The sections were then covered with mounting medium and imaged using an Axio Imager M2 (Zeiss) equipped with a 63× oil immersion objective and a Zeiss AxioCam MRc camera.

### Quantification of COX activity in *C. elegans* and cells

Prior to measuring optical density (OD), background was subtracted for all images using a “rolling ball” filter with a radius of 25 pixels for light background in FIJI (ImageJ, https://imagej.net/). Afterwards, images were inverted to negative to obtain OD values. Regions of interest were manually drawn for *C. elegans* and obtained for cell images by using the ‘create selection’ option in FIJI (ImageJ, https://imagej.net/).

### Statistical analysis

Statistical analysis was performed with GraphPad Prism (Version 10.1.1; GraphPad Software, USA). Data was tested for normal distribution by D’Agostino and Pearson omnibus normality test. Data presentation, sample size (number of biological repeats and sample size per repeat and condition), and the applied statistical tests are indicated in the Figure legends for each experiment. Statistical significance was set at an alpha level of 0.05.

## RESULTS

### Less than 0.5% of 905 nm laser light transmitted through the human skull

To assess the amount of NIR light that can reach the brain, three human skulls were illuminated with light from the 905 nm laser on the outside of the skull, while light power was measured on the inside (Figure 1A). Transmittances were measured at three distinct positions on the skulls (Figure 1B). The highest transmittance values (28-173 µW/cm^2^) were measured at the parietal position (Figure 1C). Transmittance at the frontal and occipital positions was substantially lower, with the occipital region yielding the lowest absolute power densities (3.7-30 µW/cm^2^). All three skulls showed the same transmittance pattern across positions. These variations align with reported differences in skull thickness at these sites.^25^ Measuring the incoming power of the 905 nm laser (mean value 56.6 mW/cm^2^, Figure 1D) allowed the calculation of relative transmittances (Figure 1E). Transmittances were similar with skulls 1 and 2 (on average 0.16% for skull 1 and 0.14% for skull 2), whereas skull 3 showed significantly lower transmittances (on average 0.02%) (Figure 1E). Overall, a maximum of 0.31% transmittance was observed (parietal region of skull 1), which implies that over 99.5% of the incoming laser light power was absorbed or scattered by the skulls.

**Figure 1:**
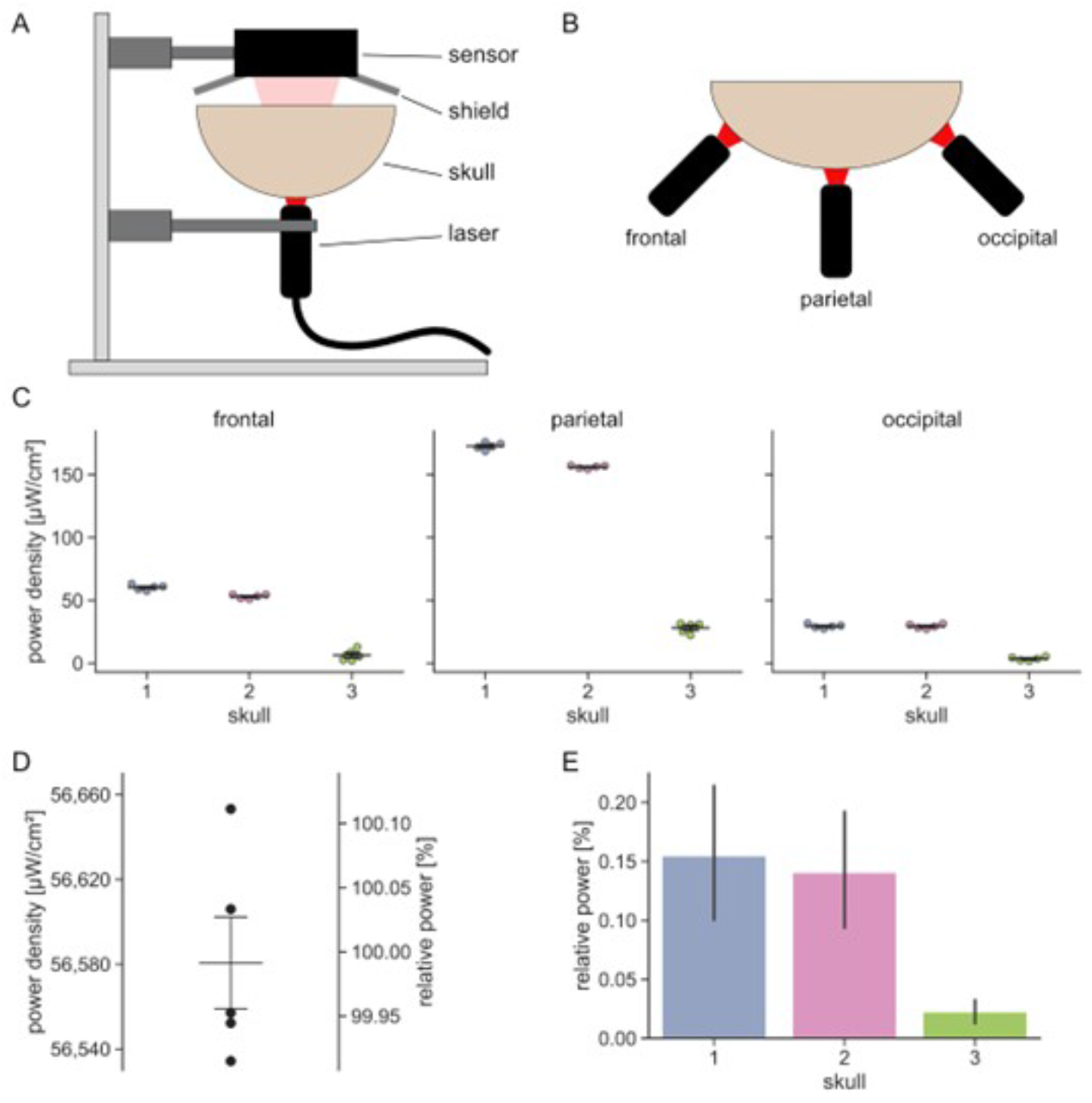
Transmittance of NIR light from the 905 nm laser through human skulls. **A** Schematic representation of the experimental setup. **B** Each skull was measured 5 times at three different positions (frontal, parietal, and occipital). **C** Absolute transmittances of NIR light measured as power density for the three skulls. Individual data points, mean and standard errors of the mean are shown. Standard errors of the mean may be difficult to identify due to their small values compared to the mean. **D** Measurements of the incoming power of the 905 nm laser in the absence of a skull. **E** Mean relative transmittances of NIR light calculated for each of the three skulls. Error bars illustrate 95% confidence intervals.

### No enhancement of mitochondrial activity in SH-SY5Y neurons and C. elegans when using 905 mm NIR light with power levels reflecting transcranial transmittances

To evaluate the effect of 905 nm NIR light exposure on brain tissue, a surrogate experimental setup was employed to simulate human *in vivo* conditions, as direct measurements in human brains are not feasible. Two experimental conditions were used: exposure to 905 nm NIR light without a filter (delivering a 134 mW/cm^2^ power density, approximating direct tissue exposure without the skull) and a filtered condition (reducing the power density to 2.5 mW/cm^2^ to mimic transcranial transmission; Figure 2A). Notably, the 2.5 mW/cm^2^ power density was still approximately 15 times higher than the 173 µW/cm^2^ maximum cranial transmittance that was measured.

**Figure 2:**
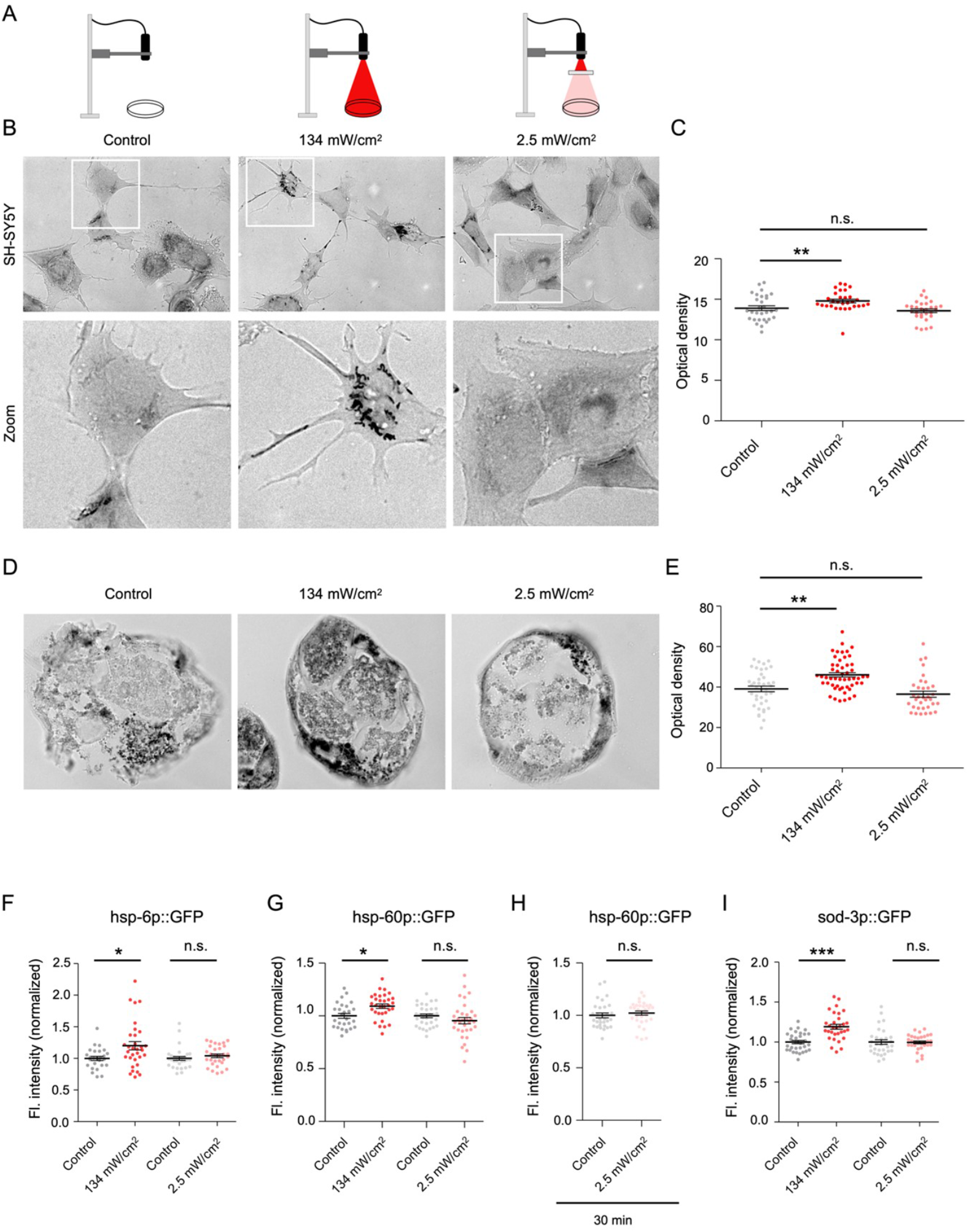
Effects of exposure to 2.5 and 134 mW/cm^2^ NIR light intensities on mitochondrial activity in SH-S5Y5 neurons and C. elegans. **A** Schematics of the experimental setup corresponding to the images in (B) and (C) and the optical density measurements in (B) and (D). **B** Representative transmitted light widefield images of SH-SY5Y neurons treated with 0 mW/cm^2^ (Control), 134 mW/cm^2^ or 2.5 mW/cm^2^ NIR light, respectively. **C** Quantification of optical density in cells treated with 0 mW/cm^2^ (Control), 134 mW/cm^2^ or 2.5 mW/cm^2^ NIR light, respectively (n=3, 10 images per condition). Individual data points, mean and standard errors of the mean are shown. Statistical analysis was performed using Kruskal-Wallis with a Dunn’s multiple comparison test. **D** Representative transmitted light widefield images of cross sections of *C. elegans* treated with 0 mW/cm^2^ (Control), 134 mW/cm^2^ or 2.5 mW/cm^2^ NIR light, respectively. **E** Quantification of optical density in cross sections of *C. elegans* treated with 0 mW/cm^2^ (Control), 134 mW/cm^2^ or 2.5 mW/cm^2^ NIR light, respectively (n=3, 10 worms per condition). Individual data points, mean and standard errors of the mean are shown. Statistical analysis was performed using a Kruskal-Wallis with Dunn’s multiple comparison test. **F-I** Quantification of fluorescence intensity (normalized to the control) of GFP expressed under the indicated promoters (*hsp-6p* and *hsp-60p*: mitochondria-specific chaperone promoters; *sod-3p*: superoxide dismutase 3 promoter) and treated with different NIR light intensities as indicated (n=27-32 from 3 independent replicates). Individual data points, mean and standard errors of the mean are shown. Statistical analysis was performed using one-way ANOVA with Bonferroni’s multiple comparison test for (F), a Kruskal-Wallis with Dunn’s multiple comparison test for (G and I), and t-test for (H). Abbreviations and symbols: n.s., not significant; *, p < 0.05; **, p < 0.01; ***, p < 0.001.

First, we assessed the impact of unfiltered 905 nm NIR light on COX activity, a known target of NIR light exposure.^4^ Exposure of SH-SY5Y neuroblastoma cells to 134 mW/cm^2^ for 5 minutes significantly increased COX substrate accumulation reflecting enhanced mitochondrial activity compared to untreated control cell**s** (Figure 2B, C). However, when the laser power density was reduced to 2.5 mW/cm^2^ the stimulatory effect on mitochondrial activity was completely abolished (Figure 2B, C). The application of NaN3, a selective COX inhibitor, completely blocked the increase in mitochondrial activity even at a power density of 134 mW/cm^2^, confirming the specificity of the observed effects (Supplementary Figure 3). These findings show that while exposure to 905 nm NIR light with a power density of 134 mW/cm^2^ stimulated mitochondrial activity, reducing NIR light power density to 2.5 mW/cm^2^ (reflecting transcranial transmission) was insufficient to provoke this effect.

To evaluate the broader biological impacts of NIR light exposure, *C. elegans* was used as an *in vivo* model. Like what was observed in SH-SY5Y cells, exposure of NIR light at an intensity of 134 mW/cm^2^ resulted in COX substrate accumulation (Figure 2D, E). Reducing the laser power density to 2.5 mW/cm^2^ eliminated the stimulatory effect on mitochondrial activity (Figure 2D, E).

To test whether higher mitochondrial activity increases the demand on the mitochondrial chaperone system, two *C. elegans* reporter strains were exposed to the 905 nm laser (134 mW/cm^2^ for 5 minutes). In a strain expressing green fluorescent protein (GFP) under the control of a mitochondria-specific chaperone promoter (*hsp-6p::GFP*), laser exposure induced chaperone expression, as indicated by elevated GFP levels (Figure 2F). Similarly, activation of the *hsp-60* promoter, regulating the expression of another mitochondria-specific chaperone, was observed (Figure 2G). However, when the laser power density was reduced to 2.5 mW/cm^2^, these effects were completely abolished (Figure 2F,G). Even extending the exposure time from 5 min to 30 min failed to induce any upregulation of chaperone expression (Figure 2H). Notably, no increased levels of the small heat shock protein HSP-16.2 were observed (Supplementary Figure 3), indicating that the 905 nm laser did not induce a heat shock response. This is consistent with previous reports that the super-pulsed design of the 905 nm laser used in this study minimizes surface heat generation^26^ and suggests a specific impact of NIR light on the mitochondrial chaperone system. Furthermore, the laser activated superoxide dismutase 3 (*sod-3p::GFP*), likely in response to increased ROS production due to enhanced mitochondrial respiration^4^ (Figure 2I). Again, this induction was not observed with the reduced power level (Figure 2I).

Taken together, while NIR light exposure at an intensity of 134 mW/cm^2^ enhanced mitochondrial activity in SH-SY5Y neurons and *C. elegans*, no such effects were observed with a NIR light power density of 2.5 mW/cm^2^. This suggests that the practical effectiveness of transcranial NIR light treatment may be constrained by its low transmittance through the skull.

### Transmittance of NIR light from LED helmets

We further investigated the transmittance of NIR light emitted from LED helmets that were specifically designed for transcranial PBM therapy. The skulls were therefore placed in two different LED helmets (810 and 1070 nm wavelength) and the sensor above the skulls measured transmitted light power (Figure 3A). Shielding was necessary to assure that only NIR light that transmitted through the skulls reached the sensor. An increasing number of LEDs was unblocked (from 1 LED to 75 LEDs) until light bypassing the skulls was detected (Figure 3B). For both NIR helmets, a notable increase in transmittance with increasing numbers of LEDs was observed (Figure 3C). To determine whether this increase was due to the higher number of unblocked LEDs or to light bypassing the skulls, incoming power of the LED helmets was measured in the absence of a skull (Figure 3D). Using these measurements, the relative transmitted power was calculated (Figure 3E). The relative transmittance data show that for higher numbers of LEDs, the proportion of NIR light that transmitted through the skull was independent of the number of LEDs used (Figure 3E). Hence, no additional NIR light that would have bypassed the skull was captured with the sensor with the experimental setup used in this study.

**Figure 3:**
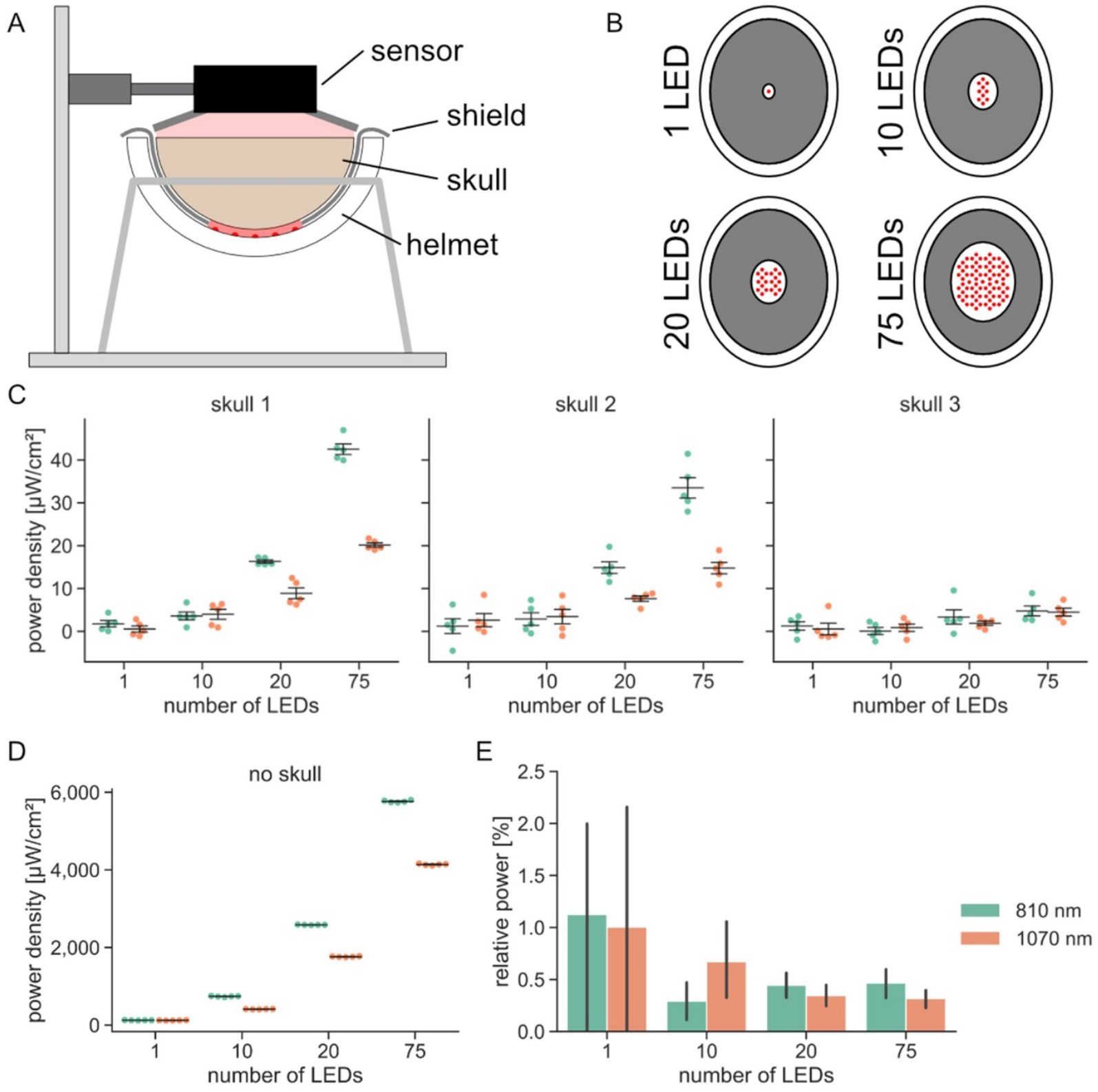
Transmittance of NIR light from two LED helmets through human skulls. **A** Experimental setup for transmittance measurements. **B** Shielding of LEDs from the helmets, required to only measure NIR light that transmitted through the skulls and avoiding NIR light travelling around the skulls. This allowed to measure transmittances with increasing numbers of LEDs. **C** Absolute transmittances of NIR light for the three skulls, measured as power density using increasing numbers of LEDs (1, 10, 20 and 75) and two different LED helmets (810 nm, green; 1070 nm, orange). Individual data points, mean, and standard errors of the mean are shown. **D** Measurements of the incoming power of the LED helmets in the absence of a skull. **E** Mean relative transmittances of NIR light calculated for each of the two LED helmets. Error bars illustrate 95% confidence intervals.

Using 1 or 10 LEDs resulted in maximum transmittances of 1.0 µW/cm^2^ (1070 nm LED helmet, skull 2) or 2.6 µW/cm^2^ (1070 nm LED helmet, skull 1) and overall high measurement errors. With the use of 20 LEDs, transmittances of up to 16.5 µW/cm^2^ (0.64%) were recorded (810 nm LED helmet, skull 1). With 75 LEDs, the 810 and 1070 nm LED helmets reached maximum transmittances of 41 µW/cm^2^ (0.71%) and 19 µW/cm^2^ (0.45%), respectively (both with skull 1). The relative power demonstrated higher spreading with the use of only 1 or 10 LEDs (Figure 3E). This was solely due to smaller measurement values when using low numbers of LEDs as the standard errors for all measurements were comparable (Figure 3C). Because the relative transmittance was highly comparable between 20 and 75 LEDs for both LED helmets, the values from the higher numbers of LEDs can be considered more reliable than the relative transmittance from 1 or 10 LEDs, which seem to be dominated by noise.

With the 810 nm LED helmet consistently higher transmittances were measured compared to the 1070 nm LED helmet. This difference was not solely attributable to the higher incoming power of the 810 nm LED helmet (Figure 3D), as it persisted with 20 and 75 LEDs even after normalizing to the incoming power (Figure 3E). Thus, the light emitted from the 810 nm LED helmet exhibited superior penetration through the skulls than light emitted from the 1070 nm LED helmet. However, the maximum transmittance measured with both LED helmets (41 µW/cm^2^ and 19 µW/cm^2^ respectively) was substantially lower than the maximum transmitted NIR light measured with the 905 nm laser (173 µW/cm^2^). Furthermore, given that these values were several orders of magnitude lower than the intensity tested in biological experiments that demonstrated no effect (2.5 mW/cm^2^), it was unnecessary to repeat these biological experiments with even lower intensities.

## DISCUSSION

This study assessed the capacity of NIR light from three commercially available devices (a 905 nm laser and two LED helmets operating at 810 and 1070 nm) to penetrate the human skull and induce a biological effect. The aim was to evaluate the potential therapeutic efficacy of transcranial PBM therapy for the treatment of neurological disorders, where NIR light is applied to the patient’s head.

Our findings reveal that more than 99% of the emitted NIR light was either absorbed or scattered by the human skull.

Depending on the skull or device used, NIR power densities between 3.7 µW/cm^2^ and 173 µW/cm^2^ were measured inside the dura mater, matching results from previous studies.^27^ Two skulls showed similar transmission values, whereas the third skull showed substantially lower transmission, a pattern observed with both the 905 nm laser and the 810/1070 nm LED helmets. Of note, the dura mater of the third skull appeared darker, possibly indicating postmortem intracranial hemorrhage, which could result in higher absorption of NIR light compared to the other skulls.

Despite these variations, the data demonstrate that less than 1% of the NIR light could penetrate the skull. While 134 mW/cm^2^ of NIR light stimulated mitochondria in cultured human neurons and *C. elegans*, a power density of 2.5 mW/cm^2^ - which was still 15 times higher than that experimentally measured - was already insufficient to trigger mitochondrial activity. This raises serious doubts as to whether the small amount of NIR light penetrating through the human skull can have any meaningful therapeutic impact.

A limitation to this study was the use of a single power sensor positioned on the other side of the human skull to measure light transmittance, which may underestimate total light penetration as it does not account for lateral scattering. However, most of the scattered light is likely absorbed by the skull and surrounding tissues before reaching brain cells, suggesting that lateral scattering would have minimal effect on the fraction of NIR light that could reach neurons in real-world conditions. In contrast, it is important to note that the use of human postmortem skulls does not directly replicate the conditions found in living patients. While shaved skin, subcutaneous tissue, cranial bone and dura mater were present, important additional light-absorbing structures such as leptomeninges and subarachnoid space containing cerebrospinal fluid and blood vessels were absent. These missing components would further reduce NIR light transmittance,^28^ not to mention that the target neurons are at a variable distances within the brain parenchyma that the light would additionally need to penetrate. Our measurements therefore overestimate the amount of light reaching cortical neurons. This overestimation is even more pronounced for deeper brain regions, such as the hippocampus, which is one of the first areas affected in AD.^29^

If this study shows that NIR light is unlikely to penetrate the skull effectively, why have some studies reported benefits? The improvements observed in patients could be related to placebo effects or the increased daily social interactions that patients experience as part of their treatment, which are known to have cognitive benefits.^30,31^ The improvements attributed to NIR therapy may therefore be due to the positive effects of patient care, attention and interaction, rather than the light therapy itself.

In conclusion, the present study provides strong evidence that NIR light from the tested devices does not penetrate the human skull in sufficient quantities to induce a photobiomodulation effect. The human skull presents a significant barrier that substantially reduces the potential efficacy of NIR treatment for brain-related diseases. Future studies should include more realistic physiological models of the human head that include structures such as leptomeninges and the subarachnoid space to fully assess the limitations of NIR light penetration *in vivo*.

## CONTRIBUTORS

Conceptualization J.T., L.K., P.R.H., C.S., C.N.K.

Formal Analysis J.T., L.K.

Investigation J.T., L.K.

Methodology J.T., L.K.

Project administration C.S., C.N.K.

Resources S.M., D.K.

Supervision C.S., C.N.K.

Visualization J.T., L.K.

Writing – original draft J.T., L.K., C.S., C.N.K.

Writing – review & editing J.T., L.K., S.M., D.K., P.R.H., C.S., C.N.K.

## DATA SHARING

The original data supporting the findings of this study, including measurements and microscopy images, as well as resources used (e.g., *C. elegans* strains), are available upon request.

## ACKNOWLEDGMENTS

We thank Beate Aschauer, Sabine Tost, Astrid Baltruschad and Barbara Mosler for excellent technical assistance. We also extend our gratitude to Claudia Harbauer, Andrea Haderer and Axel Unverzagt for preparing the human skulls. Some strains were provided by the Caenorhabditis Genetics Center, which is funded by NIH Office of Research Infrastructure Programs (P40 OD010440). In preparing this manuscript, AI technology was used solely to enhance language clarity and readability.

---

**Supplementary Figure 1:**
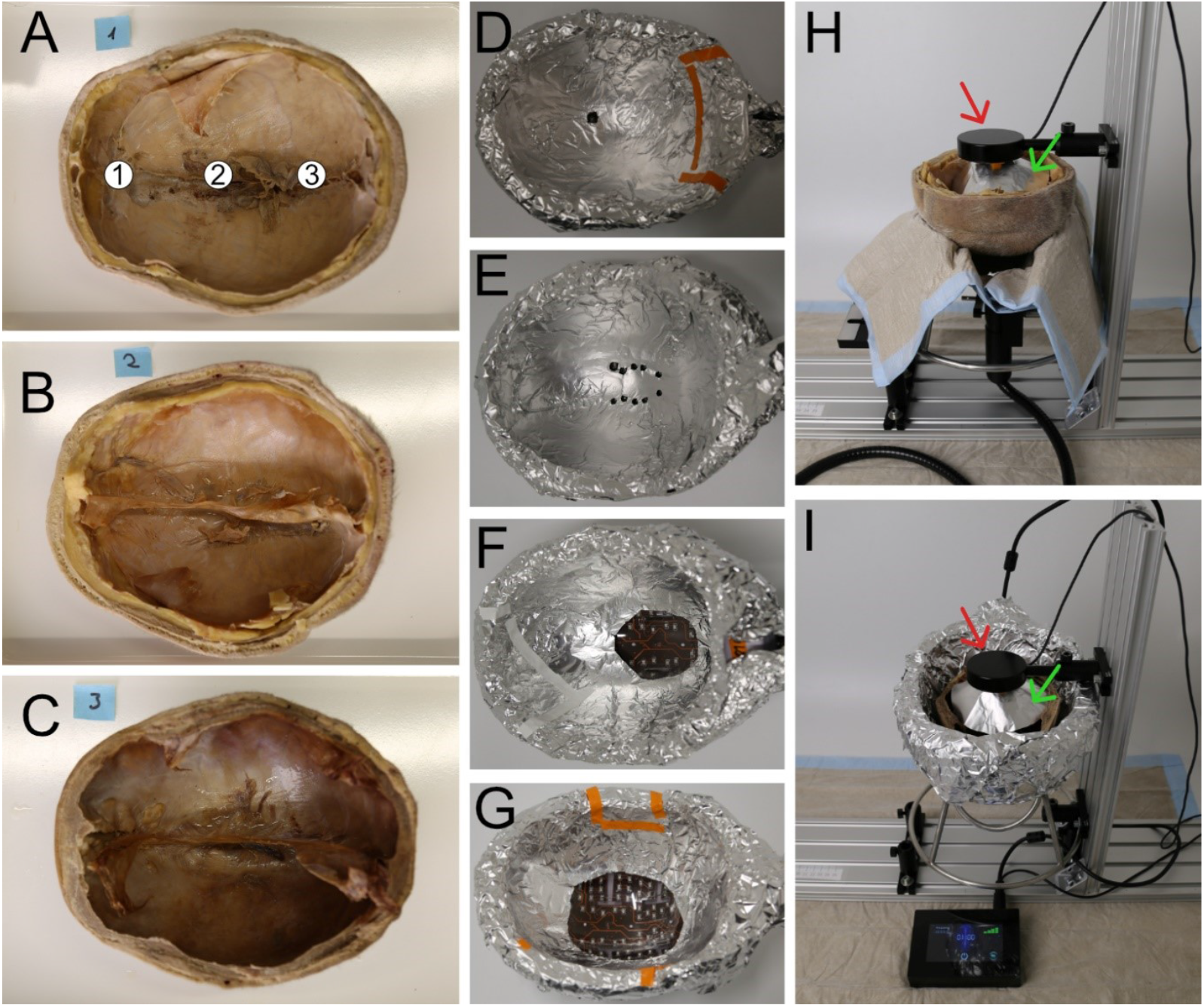
Experimental setup for transmittance measurements. **A-C** Images of the three post-mortem human skulls investigated in this study. The skulls included the skin, subcutaneous tissue, bone, and dura mater; parts of the cerebral falx were present. Using the medical laser, transmittance was measured at three different positions for each skull (marked 1-3 in panel A). **D-G** The LED helmets had 256 LEDs, each mounted on the inside of the helmets. For measurements, LEDs were partly blocked with aluminum foil allowing to measure transmittances with increasing numbers of LEDs: one LED (D), 10 LEDs (E), 20 LEDs (F), and 75 LEDs (G). **H, I** Experimental setups for the 905 nm laser (H) and the LED helmets (I). For both types of light emitters, the power sensor (red arrows) was positioned above the skulls while the emitters were positioned below. An aluminum foil shield surrounding the aperture of the power sensor (green arrows) blocked potential external light.

**Supplementary Figure 2:**
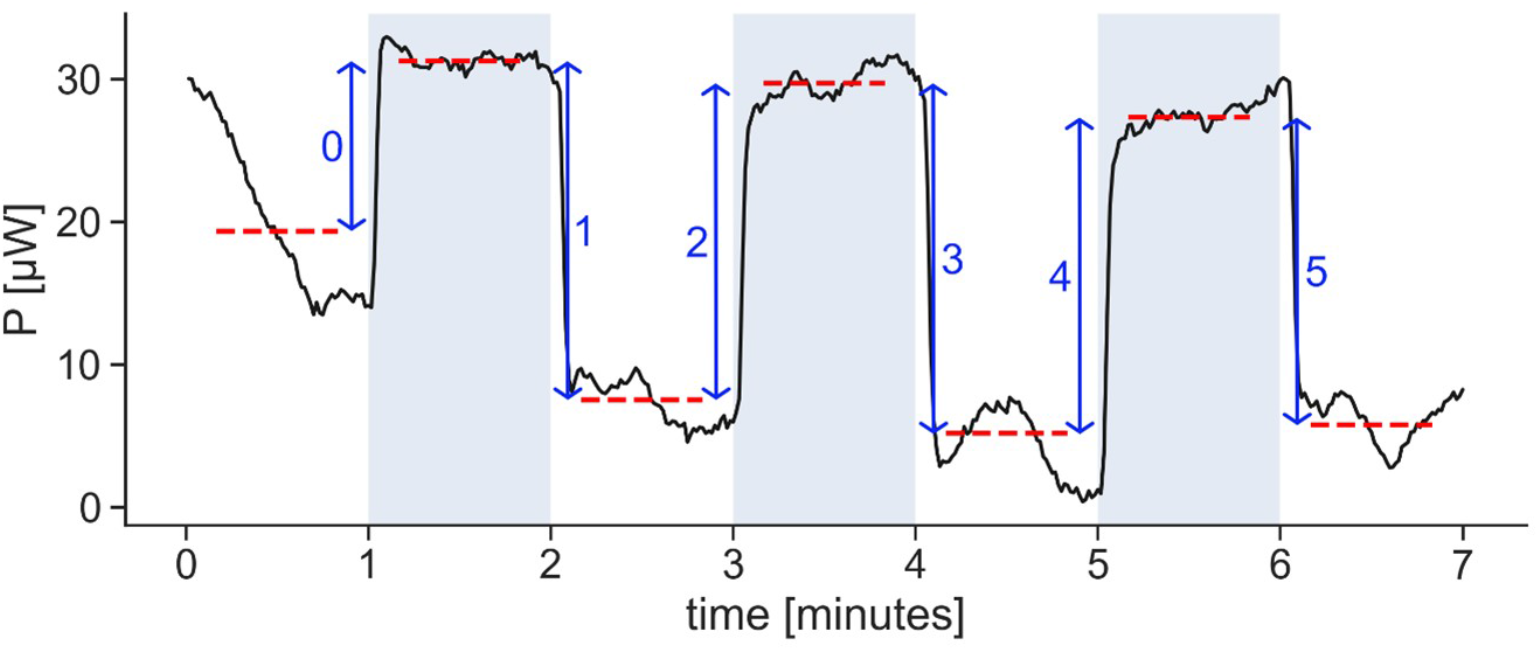
Example of raw recording (1070 nm LED helmet, 75 LEDs, skull 1) illustrating the calculation of power measurements. The recording shows the measured power (P) over time. A recording lasted for seven minutes. During this time, the light sources were turned on and off in one-minute intervals. The time during which the light source was active is depicted in blue shading. The intervals in-between were considered baseline measurements. Mean values were calculated from 40 seconds during both the active times and the baselines (red dashed lines). At each switch (off-on or on-off), a measurement was calculated as the mean on-value minus the mean baseline (blue arrows). This resulted in six measurements for each recording (labeled 0 – 5). The first measurement (0) was always discarded due to thermal instabilities and sensor drift, which can be seen in the first minute of the recording.

**Supplementary Figure 3:**
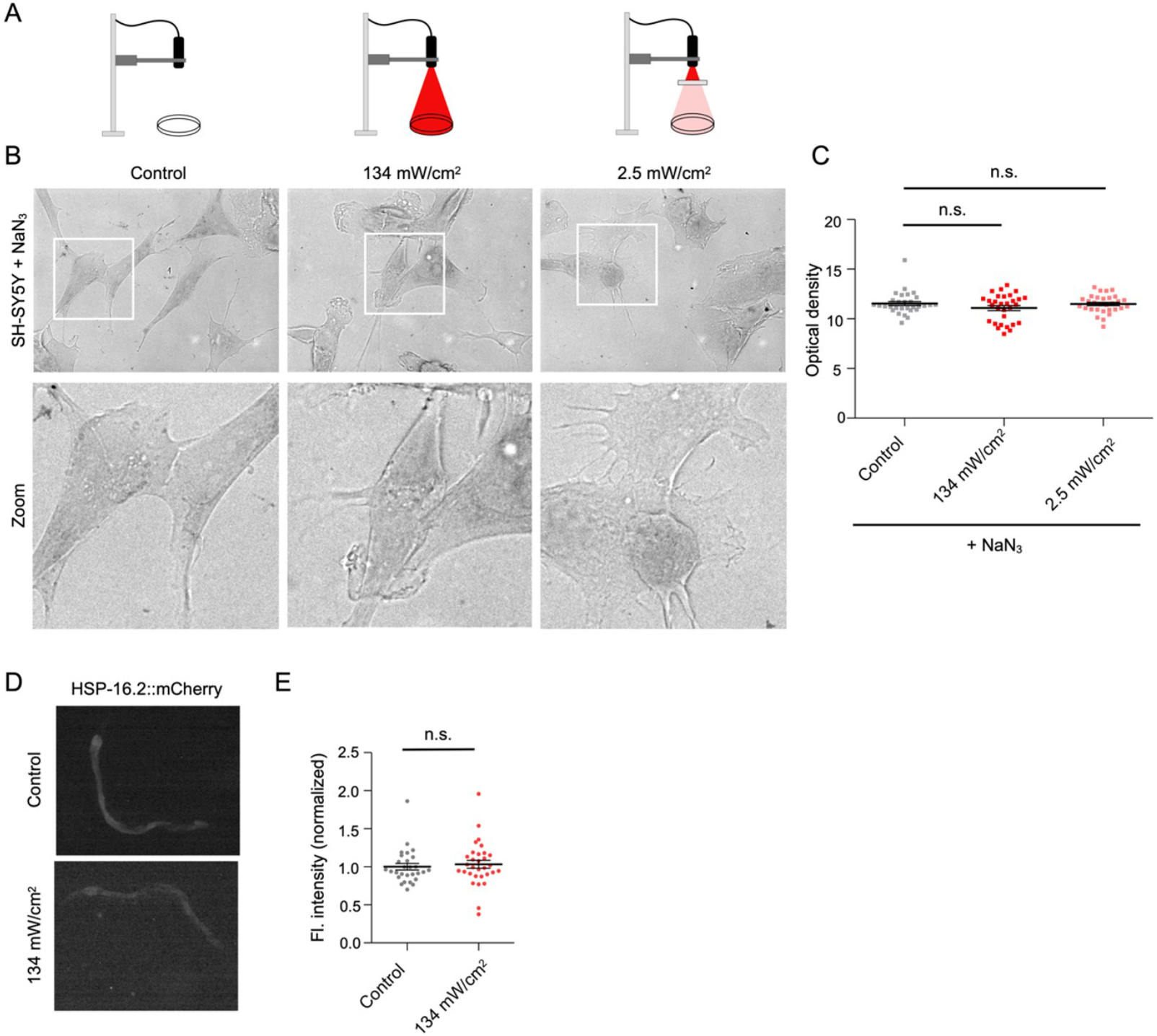
SH-SY5Y cells treated with the COX inhibitor sodium azide (NaN_3_) showed no COX activation upon NIR light treatment. **A** Schematics of the experimental setup corresponding to the images in (B) below and the optical density measurements in (C). **B** Representative transmitted light widefield images of SH-SY5Y cells treated with NaN_3_ and 0 mW/cm^2^ (Control), 134 mW/cm^2^ or 2.5 mW/cm^2^ NIR light, respectively. **C** Quantification of optical density in cells treated with NaN_3_ and 0 mW/cm^2^ (Control), 134 mW/cm^2^ or 2.5 mW/cm^2^ NIR light, respectively (n=3, 10 images per condition). Individual data points, mean and standard errors of the mean are shown. Statistical analysis was performed using a Kruskal-Wallis with a Dunn’s multiple comparison test. **D** Widefield fluorescence images of *C. elegans* expressing HSP-16.2::mCherry. **E** Quantification of HSP-16.2::mCherry fluorescence intensity (FI, normalized to the control) indicating that the heat shock response was not activated by treatment with 134 mW/cm^2^ NIR light (n=28-31 from 3 independent replicates). Individual data points, mean and standard errors of the mean are shown. Statistical analysis was done using a Mann Whitney test. Abbreviation: n.s., not significant.

